# Mechanical regulation of cell fate transitions underlying colorectal cancer metastasis formation

**DOI:** 10.1101/2023.09.17.557771

**Authors:** Mirjam C van der Net, Marjolein J Vliem, Lars JS Kemp, Carlos Perez-Gonzalez, Esther A Strating, Ana Krotenberg-Garcia, Ronja M Houtekamer, Karen B van den Anker, Jooske L Monster, Hugo JG Snippert, Antoine A Khalil, Jacco van Rheenen, Saskia JE Suijkerbuijk, Onno Kranenburg, Danijela Matic Vignjevic, Martijn Gloerich

## Abstract

Colorectal cancer (CRC) cells exhibit high plasticity and transition between different cellular states during the development of metastasis. Lgr5-expressing cancer stem cells fuel the growth of the primary tumor and metastasis, yet disseminated tumor cells arriving at the metastatic site are devoid of Lgr5 expression. It is currently unknown how CRC cell fate transitions are regulated during the metastatic process and how tumor cells give rise to metastatic lesions despite being Lgr5^neg^. Here, we show that the reprogramming of disseminating CRC cells is driven by mechanical interactions with the Collagen I-rich interstitial matrix. Collagen I-induced pulling forces are sensed by integrins and mechanosensitive calcium channels, which together direct the transition of CRC cells into a fetal-like state. The fetal-like state is maintained after reaching the blood circulation and promotes metastasis-initiation of disseminated CRC cells in the liver. Our findings indicate a key contribution of mechanical signals in controlling cell fate transitions that underlie the metastatic potential of CRC, involving an interplay between different mechanosensitive mechanisms.

## Introduction

Colorectal cancer (CRC) mortality results from its metastatic spread to distant organs, including the liver and lungs (Keum and Giovannucci, 2019). Development of metastases requires CRC cells to dynamically adapt their behavior to accomplish each step of the metastatic process. This starts with cell dissemination from the primary tumor and invasion into the underlying parenchymal tissue to reach the blood or lymph circulation. Circulating CRC cells can subsequently extravasate at distant sites and establish micro-metastatic lesions, before expanding into macro-metastases and overtaking the organ (Massagué and Obenauf, 2016; Sosa et al., 2014). Elucidation of the cellular mechanisms employed by CRC tumors to succesfully accomodate these different steps of the metastatic cascade will be instrumental to target metastasis formation.

Despite the presence of oncogenic mutations, the cellular hierarchy in CRC tumors maintains a high resemblance with the healthy intestine (Batlle and Clevers, 2017). Cancer stem cells (CSCs) marked by the expression of Lgr5 constitute a self-renewing population that gives rise to multiple cell types, with a high degree of plasticity enabling differentiated cells to regain stem cell potential (Shimokawa et al., 2017). Although Lgr5-expressing CSCs are indispensable for the growth of metastases (de Sousa e Melo et al., 2017; Fumagalli et al., 2020), recent work revealed that CRC cells disseminating into the blood circulation and arriving in the liver are predominantly devoid of Lgr5 expression (Fumagalli et al., 2020). Lgr5^pos^ CSCs reappear only after the initial stages of metastatic outgrowth (Fumagalli et al., 2020; Heinz et al., 2022), which is essential for further expansion of the tumor and development into macro-metastasis (Heinz et al., 2022). It remains unclear how CRC cell fate transitions are regulated during metastasis formation, and how Lgr5^neg^ tumor cells can give rise to initial metastatic lesions.

The malignant behavior of tumor cells is influenced by interactions with the tumor microenvironment, composed of stromal cells and the extracellular matrix (ECM). In addition to biochemical cues, tumor cells receive mechanical signals originating from physical interactions between tumor cells and their environment. For instance, confined growth of tumors induces intratumoral pressure, whereas interactions with the ECM can result in tensile forces experienced by tumor cells (Northcott et al., 2018). These forces can be sensed and transduced into intracellular biochemical signals by mechanosensors, including cell adhesion complexes, and thereby influence tumor cell behavior (Swaminathan and Gloerich, 2021). Forces experienced by the tumor can evolve during tumor progression, due to remodeling of the ECM as well as tumor cells reaching different matrix environments during the metastatic route (Montagner and Dupont, 2020). This starts with primary tumors breaching the basement membrane matrix and invading the underlying collagen I-rich interstitial matrix, which have distinct chemical and mechanical properties (Montagner and Dupont, 2020). How CRC cells respond to these biophysical changes, and whether these changes impact CRC cell fate transitions, is unclear.

Here, using CRC organoid models supported with *in vivo* CRC mouse models and histological analyses of patient tumors, we identify mechanical signals as a key regulator of the cell state transitions underlying CRC metastasis formation. We find that pulling forces between CRC cells and the fibrous collagen I network are transduced through the combined action of integrins and mechanosensitive calcium channels. These mechanosensors together direct the YAP1-mediated cellular reprogramming from a CSC into a fetal-like state, which resembles the cellular state during intestinal development and damage-induced regeneration. The fetal-like state is maintained as tumor cells circulate in the blood stream and enables the metastatic seeding and outgrowth of disseminated CRC cells. These findings indicate a key contribution of mechanical signals sensed by CRC cells, relying on integrins and mechanosensitive calcium channels, in controlling cell fate transitions that underlie their metastatic potential.

## Results

### Mechanical forces originating from tumor-collagen I interactions induce loss of Lgr5^pos^ cancer stem cells

Recent findings indicate that cells disseminating from primary CRC tumors and seeding metastases are predominantly Lgr5^neg^ (Fumagalli et al., 2020; Heinz et al., 2022). To investigate whether interactions of CRC cells with the interstitial matrix during their dissemination could be responsible for inducing loss of stemness, we mimicked these ECM alterations in a CRC organoid model. For this, we made use of organoids isolated from the tumorigenic colon of mice expressing Villin^Cre-ERT2^; APC^fl/fl^; KRAS^LSL-G12D^; TP53^KO/KO^ (A/K/P) with Lgr5^DTR/eGFP^ (Fumagalli et al., 2020). CRC organoids are typically cultured in Matrigel, which mimics the matrix composition of the basement membrane. To mimic the loss of basement membrane encapsulation and invasion into the Collagen I-rich interstitial matrix, full-grown organoids were isolated from Matrigel using dispase treatment and replated either into Collagen I gels or Matrigel (**Fig 1a**). Immunostaining confirmed that organoids transferred to Collagen I gels were devoid of a basement membrane (**Fig s1a**). When monitoring the behavior of CRC organoids embedded in these different matrices by live-cell imaging, we observed that in contrast to matrigel embedded organoids, organoids within Collagen I gels were highly dynamic with individual cells protruding in the surrounding matrix (**Fig 1b, movie 1 and 2**). Visualization of collagen fibers by reflection microscopy further showed their alignment towards the organoid, indicating that CRC cells are pulling on the collagen I network (**Fig 1c**). We next analyzed the presence of CSCs based on Lgr5^eGFP^ expression by confocal microscopy and flow cytometry, which showed that CRC organoids embedded in Matrigel contained a significant population of Lgr5^pos^ cells with clear presence of Lgr5 at the plasma membrane (**Figs 1d, s1b**). Collagen I-embedded organoids showed a strong reduction of Lgr5^pos^ CSCs, indicated by the presence of 49.1 ± 7.1 % Lgr5^low^ cells in Collagen I gels compared to 22.3 ± 10.0 % Lgr5^low^ in matrigel. (**Figs 1d, e, s1b, c**). Thus, the transition from Matrigel to Collagen I gels induces morphological changes in CRC organoids accompanied by a reduction in the number of Lgr5^pos^ CSCs.

**Figure 1.**
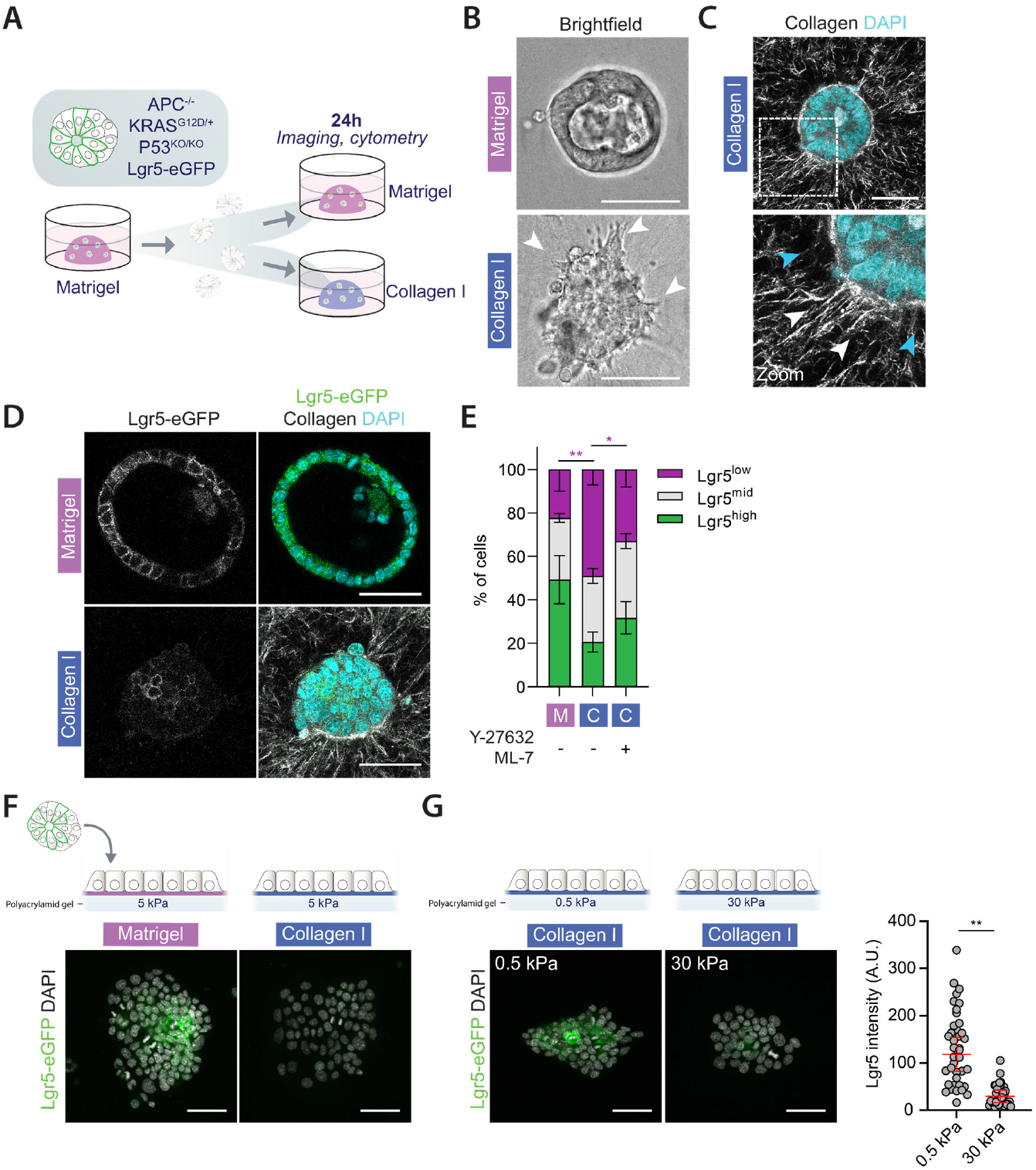
Mechanical forces originating from tumor-collagen I interactions induce loss of Lgr5^pos^ cancer stem cells. (A) Schematic overview of the embedding of organoids derived from CRC tumors in Villin^Cre-ERT2^; APC^fl/fl^; KRAS^LSL-G12D^; TP53^KO/KO^; Lgr5^DTR/eGFP^ mice in either Matrigel or Collagen I. Organoids were isolated from Matrigel by dispase treatment and subsequently replated in either Matrigel or Collagen I gels, followed by analyses after 1 day. (B) Representative examples of brightfield images from live imaging of CRC organoids embedded in Matrigel-or Collagen I gels. Arrowhead indicate protrusion of CRC cells into the Collagen I gel. Note that in collagen I gels organoids typically also lose their single-layered epithelial architecture. (C) Representative example of reflection microscopy of CRC organoid embedded in Collagen I to visualize Collagen fibers. White and blue arrows indicate aligned and non-aligned Collagen I fibers, respectively. (D) Representative images of Lgr5-eGFP expression marking cancer stem cells in CRC organoids embedded in Matrigel or Collagen I. Collagen I fibers are visualized by reflection microscopy (E) Quantification of cells with low, mid or high LGR5-eGFP expression (see Fig. s1c for further definition of expression levels) by flow cytometry of cells derived from CRC organoids embedded in Matrigel (M) or Collagen I (C), in the presence of DMSO or Y-27632 (10 μM) and ML-7 (10 μM) that was added during plating, showing mean and SD from 3 independent experiments. *P < 0.05, **p < 0.01, RM one-way ANOVA, Holm-Šídák’s multiple comparisons test comparing the Lgr5^low^ populations. (F) Representative example of Lgr5-eGFP expression in CRC organoid plated on poly-acrylamide gel (5 kPa) coated with either Matrigel or Collagen I. (G) Representative example of Lgr5-eGFP expression CRC organoid plated on poly-acrylamide gel of either 0.5 kPa or 30 kPa, coated with Collagen I. The quantification shows relative Lgr5-eGFP expression in CRC cell clusters. Data were pooled from 3 independent experiments with at least 11 clusters in each condition per experiment. Black bars represents the mean and SD of the individual experiments. **p < 0.008, ratiometric paired t-test. All scale bars represent 50 μm

Because CRC cells pull on the Collagen I network (**Figs 1b, c**), we hypothesized that these forces could act as a mechanical cue that triggers the loss of Lgr5^pos^ CSCs. Alternatively, contact with Collagen I or the loss of interactions with basement membrane components may be sufficient to induce this cell fate transition. To discriminate between these possibilities, we modulated the ability of CRC cells to exert pulling forces on the surrounding matrix, which relies on myosin-generated contractility. Inhibition of myosin activity by combined treatment with the ROCK and MLCK inhibitors Y-27632 and ML-7 demonstrated that the absence of myosin-generated forces prevented the loss of Lgr5^pos^ CSCs in Collagen I-embedded organoids (**Fig 1e**). In parallel, we tested the requirement of pulling of CRC cells on the Collagen I network by plating CRC organoids onto polyacrylamide gels functionalized with Collagen I, which recapitulated the downregulation of Lgr5^pos^ CSCs compared to organoids plated onto Matrigel-coated polyacrylamide gels (**Fig 1f**). The level of pulling forces on the Collagen I matrix can be modulated by adjusting the stiffnesses of the polyacrylamide gels (Ghibaudo et al., 2008). Only on stiff substrates on which cells are able to exert strong pulling forces (≥ 5kPa) we observed the Collagen I-induced downregulation of Lgr5^pos^ CSCs (**Figs 1f, 1g**), whereas this reduction was not observed on soft (0.5 kPa) Collagen I-coated gels (**Fig 1g**). These findings demonstrate that instead of merely contact with Collagen I (or loss of basement-membrane contact), the ability to pull on Collagen I fibers is required for the loss of Lgr5^pos^ CSCs in CRC organoids.

### Collagen I interactions drive the transition of CRC cells into a fetal-like state

Having shown that mechanical interactions with Collagen I induce the loss of CSCs, we next determined into which state CRC cells in Collagen I-embedded organoids have transitioned. To identify different cell populations, we performed single-cell RNA sequencing of organoids derived from either Matrigel or Collagen I matrices (**Fig 2a**). Comparison of organoids from these conditions showed a separation of their transcriptional landscapes, with 5 individual cell clusters being identified when applying unsupervised clustering (clusters 0-4, **Fig 2b**). Two of these clusters (clusters 2 and 3) are characterized by high expression of intestinal stem cell markers (e.g. *Lgr5, Ascl2 and Axin2*) and originate predominantly from Matrigel-embedded organoids (**Figs 2c-2e**). Cells from Collagen I-embedded organoids are mainly present in two other clusters (clusters 0 and 4), characterized by a low abundance of Lgr5 and other classical stem cell markers (**Fig 2c-e**). Differential gene expression analysis revealed genes related to the actin cytoskeleton, migration and glycolysis to be upregulated in cells derived from Collagen I-embedded cultures (**Fig s2b**). In addition, the transcriptional profile of Collagen I-derived organoids strongly overlapped with genes expressed in spheroids isolated from the fetal intestine (**Figs 2f, s2c**) (Mustata et al., 2013). Genes that were most prominently upregulated in Collagen I-embedded organoids were associated with this fetal signature, including the *bona fide* fetal-state markers *Ly6a/Sca-1* and *Anxa1* (**Figs 2h, s2a)**. In line with this, Collagen I-embedded organoids showed upregulation of genes linked to regeneration following Dextran Sulfate Sodium (DSS)-induced Colitis, which has previously been shown to be associated with a transition of cells into a fetal-like state (**Fig 2g, s2c**) (Yui et al., 2017). We validated the observed transition of CRC cells interacting with Collagen I into a fetal-like program by immunostainings analyzed by confocal microscopy and flow cytometry, showing that the downregulation of Lgr5-eGFP is accompanied by upregulation of markers of the fetal-like state; Sca-1 and AnnexinA1 (encoded by *Anxa1*) (**Figs 2i**,**j** and **s2d**,**e**). The Collagen I-induced cell fate transition is not restricted to the murine CRC organoid model, as human organoids derived from CRC tumors (**Fig s2f**) and non-malignant mouse small intestinal organoids (Yui et al., 2017; Ramadan et al., 2022) underwent a similar upregulation of fetal makers following Collagen I interactions.

**Figure 2.**
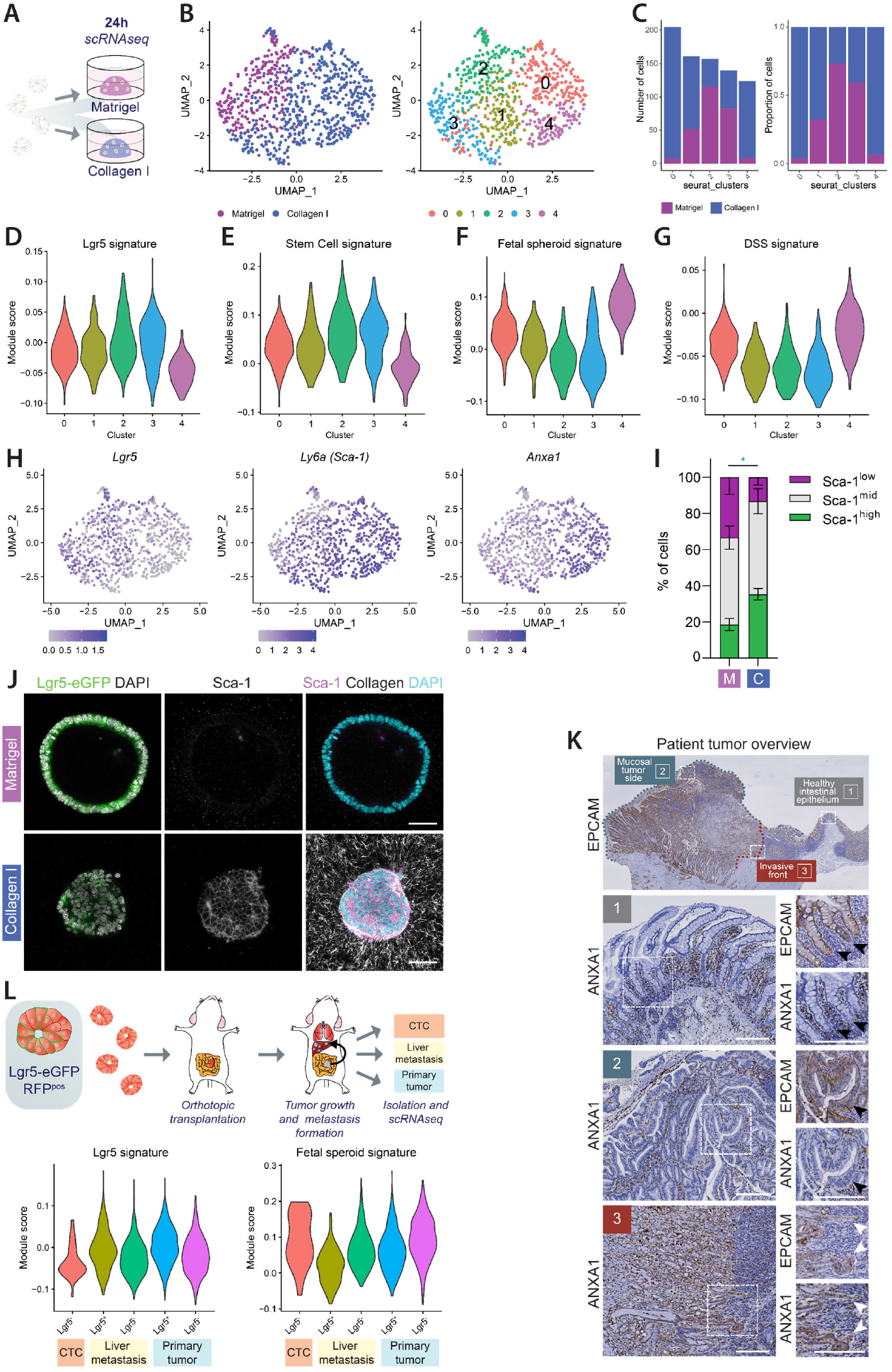
Collagen I interactions drives the transition of CRC cells into a fetal-like state. (A) Schematic overview of the single cell RNA sequencing analysis of CRC organoids 1 day after embedding in either Matrigel or Collagen I. (B) Reduced dimensionality (UMAP) visualization of single cell RNA sequencing data of cells originating from Matrigel-or Collagen I embedded CRC organoids, with color-coding by matrix origin (left) and by results of unsupervised clustering (right). (C) Composition of clusters color-coded by matrix origin, with absolute number (left) and proportion (right) of cells. (D) Violin plot showing single cell module scores for expression of genes from the Lgr5 cell signature (Muñoz et al., 2012). (E) Violin plot showing single cell module scores for expression of genes from the intestinal stem cell signature (Haber et al., 2017). (F) Violin plot showing single cell module scores for expression of genes from the fetal spheroid signature (Mustata et al., 2013). (G) Violin plot showing single cell module scores for expression of genes from the DSS-induced colitis signature (Yui et al., 2017). (H) Reduced dimensionality (UMAP) visualization with color-coded gene expression levels of Lgr5, Ly6a (Sca-1) and Anxa1. (I) Quantification of cells with low, mid or high Sca-1 expression by flow cytometry (see Fig. s2d for further definition of expression levels) of cells derived from CRC organoids embedded in Matrigel (M) or Collagen I (C), showing mean ± SD from 3 independent experiments. *P < 0.05, paired t-test comparing the Sca-1^high^ populations. (J) Representative images of Lgr5-eGFP expression and Sca-1 immunostaining in CRC organoids embedded in Matrigel or Collagen I. Collagen I fibers are visualized with reflection microscopy. Scale bar represents 50 μm. (K) IHC stainings for the epithelial marker EPCAM and fetal-state marker AnnexinA1 (ANXA1) of an invasive CRC tumor, comparing the healthy intestinal epithelium, non-invasive mucosal side, and invasive tumor front. Arrows indicate EPCAMpos tumor regions with either absence (black arrow) or presence (white arrow) of ANXA1 expression. Note that ANXA1 is also expressed in EPCAM^neg^ stromal cells. Scale bar represents 200 μm. (L) Analysis of single cell RNA sequencing (Fumagalli et al., 2020) of cells isolated from the circulation (CTCs), primary tumor or liver metastasis after orthotopic transplantation of APC^fl/fl^; KRAS^LSL-G12D^; TP53^KO/KO^ (A/K/P) Lgr5^DTR/eGFP^ organoids in mice. Violin plots show single cell module scores for expression of genes from an Lgr5 cell signature (Left, (Muñoz et al., 2012)) and a fetal spheroid signature (Right, (Mustata et al., 2013)).

Because our organoid model showed that Collagen I interactions induce the transition of CRC cells into a fetal-like state, we next analyzed the expression of fetal-state markers during the invasion of tumors into the Collagen I-rich interstitial stroma in patients. Histological analyses for the fetal state marker AnnexinA1 (Vasquez et al., 2022) showed a clear presence of AnnexinA1^pos^ cells in primary tumors of CRC patients (**Fig 2k**). These AnnexinA1^pos^ cells were present at the invasive tumor margins, particularly in tumor regions associated with the surrounding stroma (**Fig 2k**). In contrast, cells in the tumor bulk or healthy epithelium showed very low to no expression of AnnexinA1 (**Fig 2k**). To test whether cells that eventually disseminate from primary CRC tumors also reside in this state, we analyzed the transcriptional profile of CRC cells that have entered the blood circulation in a colonic orthotopic transplantation mouse model (Fumagalli et al., 2020). Circulating tumor cells were isolated from blood of the portal vein of mice bearing metastatic CRC, which were previously shown to lack classical stem cell markers (Fumagalli et al., 2020). We then performed more detailed transcriptional analysis of these circulating tumor cells and compared them to Lgr5^neg^ and Lgr5^pos^ cells isolated from the primary tumor and liver metastasis, which showed that circulating tumor cells display elevated expression of a fetal-like gene signature (**Fig 2l**). Altogether, our findings imply that CRC cells transition into a fetal-like state due to interactions of disseminating CRC tumor cells with the Collagen I-rich interstitial matrix which is maintained when these cells reach the metastatic site.

### Transition into a fetal-like state promotes outgrowth of metastatic lesions

Previous studies in the healthy intestine demonstrated that the transition of intestinal cells into a fetal-like state drives the proliferative response during regeneration of the intestinal epithelium after several types of damage (Yui et al., 2017; Ayyaz et al., 2019; Nusse et al., 2018). Akin to this regenerative response, the fetal-like state of CRC cells may enable metastatic seeding and outgrowth of disseminated Lgr5^neg^ cells at distant sites. To test whether Lgr5^neg^ cells are able to successfully grow out due to their transition into a fetal-like state, we isolated single Lgr5^low^ cells and classified them on the presence or absence of a fetal-like gene signature based on expression of Sca-1. We then followed the outgrowth of these cells into multicellular organoids *in vitro* over time (**Fig 3a**). Sca-1^high^ cells showed a strong increase in outgrowth efficiency compared to Sca-1^low^ cells (40.6% versus 14.5% at T=40 hours, respectively, **Figs 3b**,**c**). Furthermore, the organoids formed from Sca-1^low^ cells were significantly smaller than those formed from Sca-1^high^ cells (**Fig 3d**).

**Figure 3.**
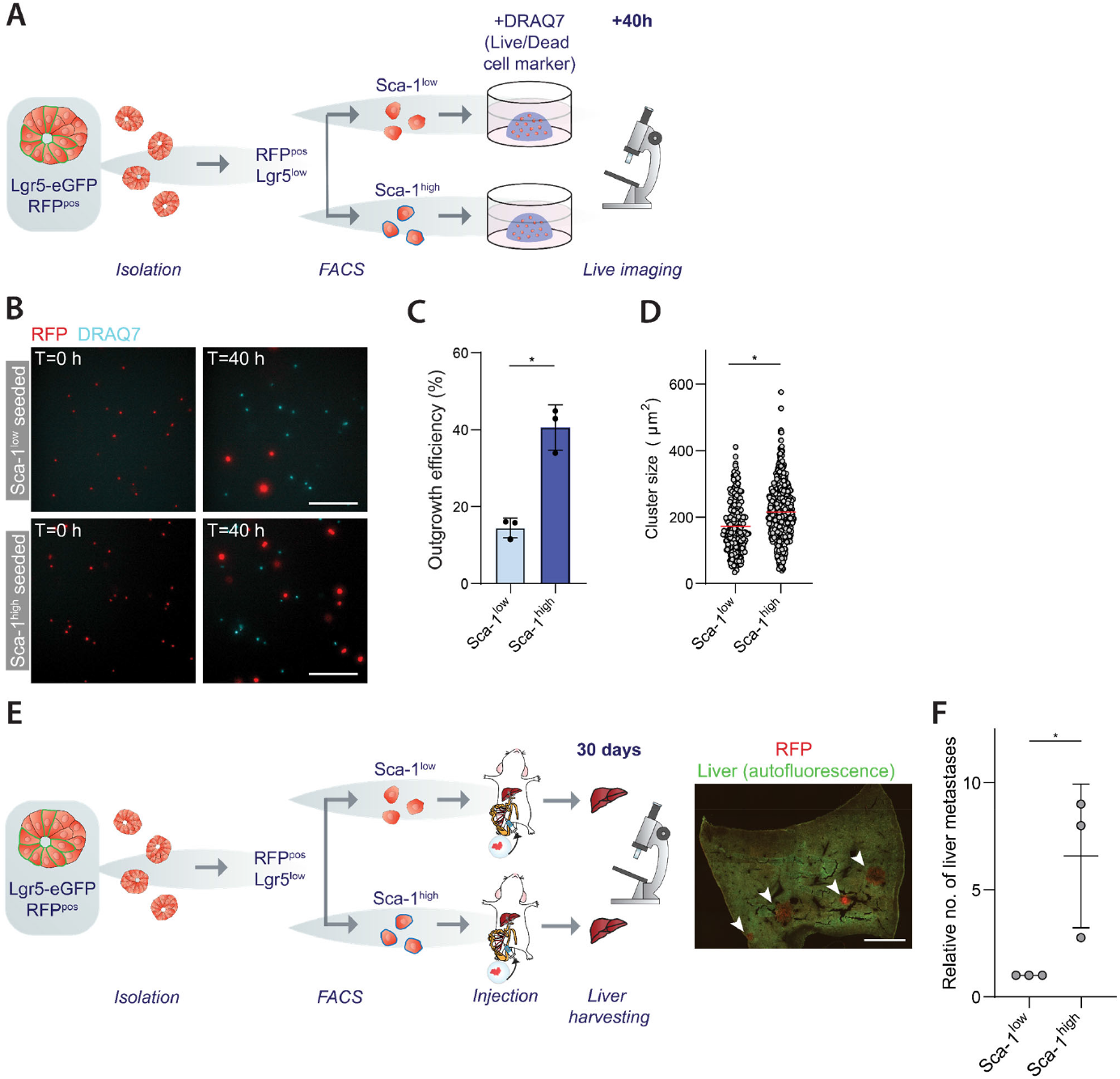
Transition into a fetal-like state promotes outgrowth of metastatic lesions. (A) Schematic overview of the experimental setup of the *in vitro* outgrowth of single Lgr5^low^ cells with either high or low levels of Sca-1 expression. Single cells being RFP^pos^, Lgr5^low^ and DRAQ7^neg^ were isolated by FACS and separately collected based on the expression of Sca-1. Isolated cells were seeded in Collagen I and followed with time-lapse imaging, with dead cells being marked by DRAQ7. (B) Representative images of time lapse microscopy following the survival and outgrowth of Sca-1^low^ and Sca-1^high^ single cells in Collagen I. Scalebar represents 200 μm. (C) Quantification of the outgrowth efficiency of Sca-1^low^ and Sca-1^high^ single cells 40 hours following their seeding, showing mean ± SD outgrowth % from three independent experiments. *P < 0.05; paired t-test. (D) Quantification of the cluster size (μm^2^) of surviving Sca-1^low^ and Sca-1^high^ single cells 40 hours following their seeding, showing grand median size. Data was pooled from three independent experiments, with every dot representing a cell cluster. *P < 0.05; paired t-test of the means of the independent experiments. (E) (Left) Schematic overview of the experimental setup of the metastatic outgrowth of single Lgr5^low^ cells with either high or low levels of Sca-1 expression injected in the mesenteric vein of recipient mice. RFP^pos^ Lgr5^low^ DRAQ7^neg^ cells were isolated by FACS and separately collected based on the expression of Sca-1 and injected as equal numbers in the mesenteric vein. After 4 weeks the livers were harvested and subjected to analysis with microscopy. (Right) Representative image of a (30 μm thick) liver slice showing RFP^pos^ metastatic foci. Scalebar represents 2 mm. (F) Relative number of metastases per liver originating from Sca-1^low^ or Sca-1^high^ cells injected in the mesenteric vein, normalized to number of metastasis in livers from Sca-1^low^-seeded cells of each experiment with independently performed sorting and surgeries. Mean ± SD from 3 separate mice per group. *P < 0.05; ratio paired t-test.

As our experiments indicate that the fetal-like state promotes outgrowth potential *in vitro*, we next determined whether this cellular state is needed for circulating tumor cells that reside in a Lgr5^neg^ state to develop liver metastases. For this, we isolated single Lgr5^low^ cells with either low or high Sca-1 expression and injected these cells into the mesenteric vein of mice, which directly drains to the liver (**Fig 3e**). Analysis of metastases in the liver 4 weeks post-injection showed a significant increase in metastases formed in the livers of mice injected with Sca-1^high^ compared to Sca-1^low^ cells (**Fig 3f**). We often noted the presence of Sca-1^pos^ cells in those metastatic foci that could develop from Sca-1^low^ cells (**Fig s3a**). Furthermore, all outgrowing metastases showed the presence of Lgr5^pos^ CSCs (**Fig s3b**), thus underscoring the plastic behavior of these cells. These data show that the fetal-like state promotes the outgrowth capacity of CRC cells and explains the ability of disseminating Lgr5^neg^ cells to develop metastases.

### The Collagen I-induced transition into a fetal-like state is mediated by YAP1

The transcriptional transactivator YAP1 has previously been implicated in the reprogramming of healthy intestinal cells into the fetal-like state (Yui et al., 2017; Guillermin et al., 2021). In line with previous reports, our scRNAseq analysis of CRC organoids interacting with Collagen I indicated upregulation of a YAP1-dependent transcriptional signature compared to Matrigel-embedded organoids (**Fig s2c**). This was further confirmed by analyzing transcription targets of YAP1 (*Ankrd1, Ctgf*) by RT-qPCR (**Fig 4d**). In line with this induction of YAP1-mediated transcription, analyses of the cellular distribution of YAP1 by confocal microscropy revealed increased nuclear accumulation of YAP1 in CRC organoids interacting with Collagen I compared to Matrigel-embedded organoids (**Figs 4a, b**).This nuclear enrichment of YAP1 in Collagen I gels was most pronounced in cells localized at the outside of organoids (**Fig 4a, c**), and often particularly prominent in cells in contact with aligned fibers of the Collagen I network (**Fig 4a**).

**Figure 4.**
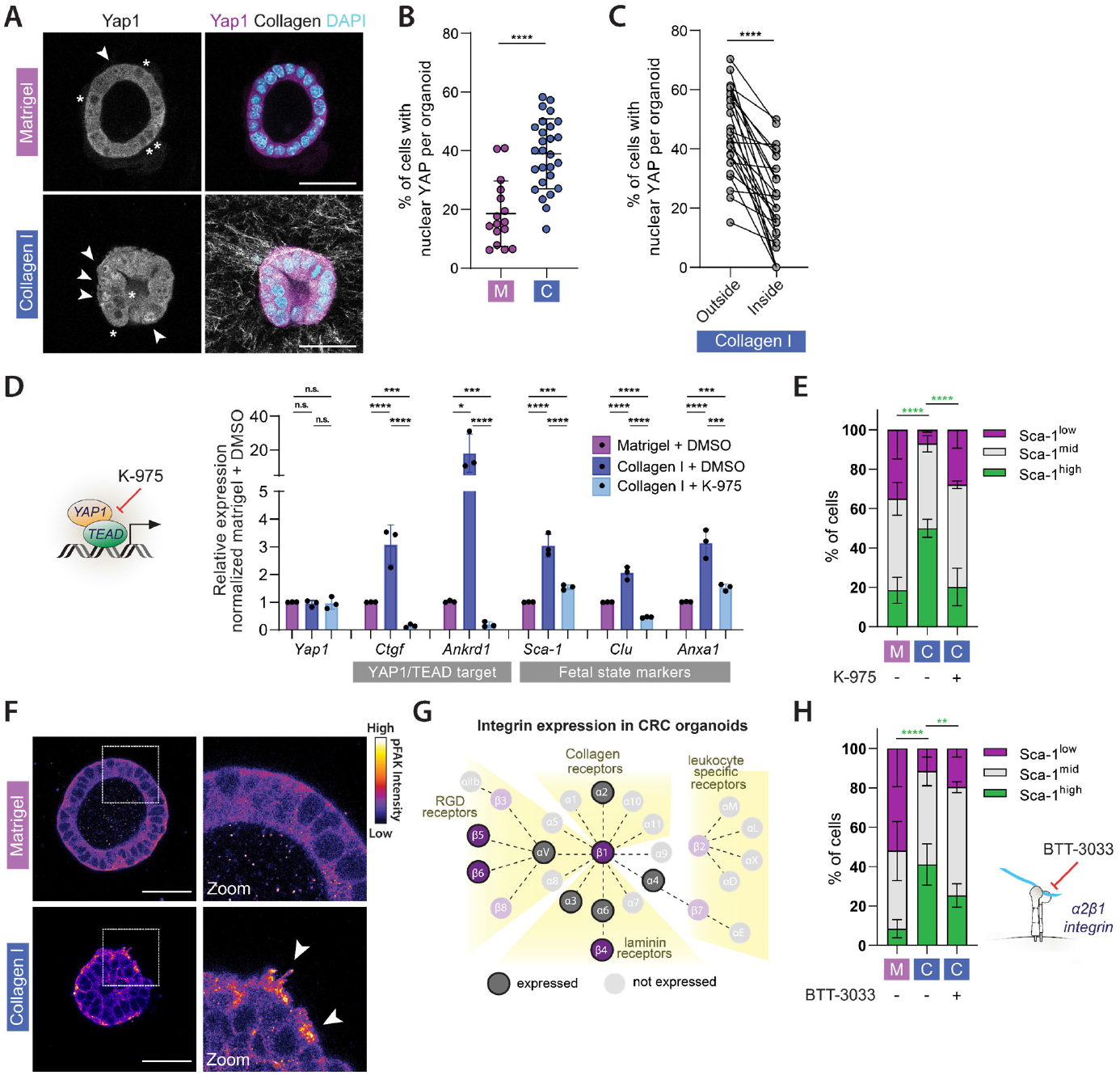
The Collagen I-induced transition into a fetal-like state is mediated by YAP1. (A) Representative images of Yap1 immunostainings of CRC organoids embedded in Matrigel or Collagen I. Collagen I fibers are visualized with reflection microscopy. Arrows indicate cells with nuclear YAP1, asterics indicate cells with nuclear-excluded YAP1. Scale bar represents 50 μm. (B) Quantification of the percentage of cells per organoid with nuclear Yap1 enrichment in organoids embedded in Matrigel (M) or Collagen I (C). Every data point represent a single organoid, data were pooled from 2 independent experiments with at least 8 organoids per condition in each experiment. Black bars represent mean ± SD. ***P < 0.005, unpaired t test. (C) Quantification of the percentage of cells with nuclear Yap1 per organoid in CRC organoids embedded in Collagen I, comparing cells localized in the outer rim and inside of the organoid. Every data point represent a single organoid, data were pooled from 2 independent experiments with 10 organoids per condition in each experiment. Black bars represent mean ± SD. ***P < 0.005, unpaired t test. (D) (Left) Schematic representation of K-975 targeting the YAP1/TEAD transcriptional complex by blocking the binding of YAP1 and TEAD. (Right) RT-qPCR analysis of CRC organoids embedded in Matrigel or Collagen I, in the presence of DMSO or the YAP1/TEAD inhibitor K-975 (10 μM) that were added during plating, showing mean expression ± SD from 3 independent experiments. *p < 0.05, **p < 0.01, ***p < 0.005, ****p < 0.001; RM-one-way ANOVA with Holm-Šídák’s multiple comparisons test performed comparing the ΔΔCT values per gene. (E) Quantification of cells with low, mid or high Sca-1 expression by flow cytometry of cells derived from CRC organoids embedded in Matrigel (M) or Collagen I (C), either in presence of DMSO or the YAP1/TEAD inhibitor K-975 (10 μM) that were added during plating, showing mean % ± SD from 3 independent experiments. ***P < 0.005, **** P < 0.001, RM-one-way ANOVA with Holm-Šídák’s multiple comparisons test comparing the Sca-1^high^ populations. (F) Representative images of CRC organoids embedded in Matrigel or Collagen I and immunostained for FAK pY397. Scale bar represents 50 μm. (G) Schematic overview of the various pairs of integrin subunits (indicated with dotted lines) and their ligands, indicating which subunits are expressed (shown non-transparent) in CRC organoids based on scRNA sequencing analysis (Fig. 2A). (H) (Left) Quantification of cells with low, mid or high Sca-1 expression by flow cytometry of cells derived from CRC organoids embedded in Matrigel (M) or Collagen I (C), either in presence of DMSO or the α2β1-integrin inhibitor BT-3033 (5 μM) that were added during plating, showing mean % ± SD from 3 independent experiments. **P < 0.01, **** P < 0.001, RM-one-way ANOVA with Holm-Šídák’s multiple comparisons test comparing the Sca-1^high^ populations. (Right) Schematic representation of BTT-3033 blocking the interaction of α2β1-integrin with Collagen.

To test whether YAP1 transcriptional activity is essential for the Collagen I-induced transition of CRC cells into a fetal-like state, we inhibited YAP1-dependent transcription using K-975, which selectively blocks the interaction of YAP1 with its partnering transcription factor TEAD (Kaneda et al., 2020) (**Fig 4d**). qPCR analyses demonstrated that upon Collagen I interaction, the upregulation of YAP1 transcriptional targets as well as fetal genes were significantly attenuated in the presence of K-975 (**Fig 4d**). In line with this, inhibiting YAP1 transcriptional activity in Collagen I-plated organoids inhibited the loss of Lgr5^pos^ stem cells and the increase of Sca-1^high^ cells (**Figs 4e, s4a**). Conversely, expression of a mutant of YAP1 that consititutively resides in nucleus (YAP1-5SA; in which residues mediating its cytosolic sequestration are mutated), was sufficient to induce the transition into a fetal-like state of CRC organoids embedded in Matrigel (**Figs S4b**,**c**). Altogether, these data demonstrate that induction of YAP1-dependent transcription drives the fate switch of CRC cells towards a fetal-like state.

Having established that mechanical interactions with Collagen I induces YAP1 transcriptional activation to drive the reprogramming of CRC cells, we set out to investigate how CRC cells relay this mechanical information to YAP1. We first determined the contribution of integrin receptors, which directly interact with the collagen network and are implicated in transducing mechanical cues from the ECM to the regulation of YAP1 signaling (Dupont, 2016). We analyzed the activity status of Focal Adhesion Kinase (FAK), the central component in the transduction of integrin-ECM interactions to intracellular signaling events (Sun et al., 2016). Immunostaining for active FAK (pY397) showed elevated FAK activity in Collagen I-embedded CRC organoids compared to their counterparts in Matrigel, particularly at regions in contact with strongly aligned collagen I fibers (**Figs 4f** and **s4d**). Our scRNAseq analysis revealed that CRC organoids express one Collagen I-interacting integrin subtype, which is α2β1-integrin (**Fig 4g**). Functionally blocking of this α2β1-integrin by BTT-3033 significantly reduced the emergence of Sca-1^high^ cells in Collagen I gels (**Fig 4h**). Altogether, our data indicate a role for integrin and downstream YAP1 signaling in Collagen I-induced reprogramming of CRC cells into a fetal-like state.

### Integrins and mechanosensitive calcium channels coordinate the Collagen I-induced fetal-state transition

As the YAP1-mediated induction of the fetal-like state was not fully inhibited upon blocking signaling through Collagen I-binding integrins (**Fig 4h**), this raises the possibility that other mechanosensors may contribute to the transduction of collagen I-associated forces to regulate CRC cell fate. Because recent findings in fibroblasts revealed that mechanosensitive calcium channels of the Piezo-and Trp families are activated at sites of pulling of cells on the ECM (Pardo-Pastor et al., 2018; Ellefsen et al., 2019), we tested whether these mechanosensors participate in the mechanical regulation of CRC reprogramming. To monitor intracellular calcium dynamics we generated CRC organoids stably expressing the genetically encoded K-GECO-1 calcium sensor (**Fig 5a**) (Shen et al., 2018). Live-imaging of this sensor in organoids embedded in Matrigel revealed minimal calcium dynamics with cells only sporadically showing calcium spikes (average frequency of 0-0.1 calcium peaks / min in individual cells) (**Fig 5 a-c, movie 3**). In strong contrast, CRC organoids in Collagen I showed strong fluctuations of intracellular calcium levels, with significant increases in both the frequency (average frequency of 0-0.3 calcium peaks / min in individual cells) and amplitude of calcium spikes (**Fig. 5 a-d, movie 4**). These enhanced calcium influxes were particularly pronounced in cells closely associated with aligned collagen I fibers (**Fig 5e**), suggesting that collagen I pulling results in opening of mechanosensitive calcium channels. We therefore tested whether activation of these channels is required for the Collagen I-induced CRC cell state transition through incubation with the spider venom peptide GsMTx4 (Suchyna et al., 2000; Gnanasambandam et al., 2017). This peptide, which targets mechanosensitive channels of the Piezo-and Trp-families, reduced the emergence of Sca-1^high^ cells in Collagen I-embedded organoids (**Fig 5f, g**). Whereas individual stimulation with the α2β1-integrin inhibitor BTT-3033 and GsMTx4 both showed a partial inhibition of the reprogramming of CRCs (**Fig 5g**), their combination resulted in near complete block of the Collagen I-induced transition of cells into the fetal-like state (**Fig 5g)**. Finally, we tested whether opening of mechanosensitive calcium channels in the absence of Collagen I-associated pulling forces can induce CRC cell reprogramming. For this, we treated organoids embedded in Matrigel with Yoda1, a chemical activator of the mechanosensitive calcium channel Piezo (**Fig 5f**). Yoda1-mediated Piezo activation induced YAP1-mediated transcription, as indicated by the upregulation of YAP1 target genes (**Fig 5h**). Moreover, direct Piezo activation was sufficient to drive the reprogramming of Matrigel-embedded organoids into the fetal-like state (**Figs 5i**,**j**), which was dependent on YAP1-TEAD transcriptional activity (**Fig 5i**). These data demonstrate that integrin and mechanosensitive calcium channels together coordinate the YAP1-dependent reprogramming of CRC cells following mechanical interactions with the Collagen I network. Altogether, our findings indicate that mechanical interactions of disseminating CRC cells with the surrounding matrix are relayed by different mechanosensors to reprogram CRC cells into a fetal-like state, which promotes their metastatic outgrowth when reaching target organs.

**Figure 5.**
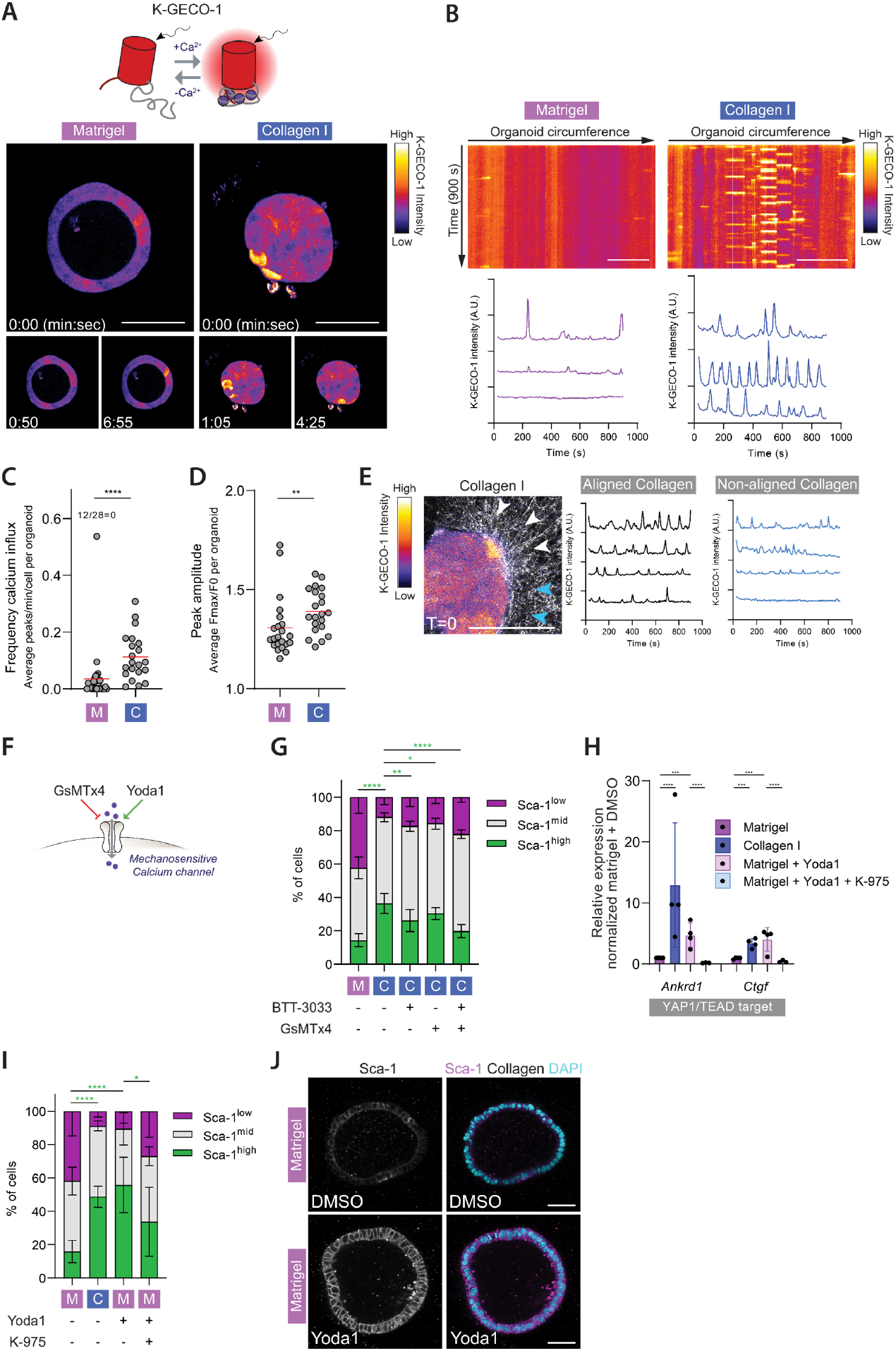
Integrins and mechanosensitive calcium channels coordinate the Collagen I-induced fetal-state transition. (A) Still images from time-lapse microscopy of CRC organoids expressing the K-GECO-1 calcium sensor (Shen et al., 2018). (B) Representative examples of calcium dynamics in Matrigel-and Collagen I-embedded CRC organoids imaged with the K-GECO-1 calcium sensor. Calcium dynamics over time in cells in the organoid circumference in contact with the surrounding matrix are shown in kymographs (top) and representative traces (bottom). Scale bar represents 50 μm. (C) Quantification of the average calcium peak frequency per minute per cell in individual organoids, imaged with the K-GECO-1 calcium sensor. Every data point represent a single organoid, data were pooled from 2 independent experiments with at least 10 organoids per condition in each experiment. Red bars represent mean. ****P < 0.001, Mann Whitney test. (D) Quantification of average peak fold change from baseline calculated by Fmax (the peak maximal intensity) / F0 (baseline intensity). Every data point represent a single organoid, data were pooled from 2 independent experiments with at least 10 organoids per condition in each experiment. Red bars represent mean. **P < 0.01, Mann Whitney test. (E) Representative example of calcium traces in CRC organoids embedded in Collagen I imaged with the K-GECO-1 calcium sensor, comparing cells in contact with aligned-or non-aligned Collagen I fibers. Left: Still image of time-lapse microscopy of the K-GECO-1 sensor (Fire) with Collagen I (white). Scalebar represents 50 μm. Arrows indicate aligned (white) and non-aligned (blue) Collagen I fibers. Right: Representative single-cell traces of the relative K-GECO-1 intensity of cells in contact with aligned (left) or non-aligned (right) Collagen I fibers. (F) Schematic representation of the action of GsMTx4 and Yoda1. GsMTx4 inhibits mechanosensitive calcium channels of the Piezo-and TRP families, and Yoda1 reduces the threshold of membrane tension for Piezo1 channel opening. (G) Quantification of cells with low, mid or high Sca-1 expression by flow cytometry of cells derived from CRC organoids embedded in Matrigel (M) or Collagen I (C), either in presence of DMSO, the α2β1-integrin inhibitor BT-3033 (5 μM), the mechanosensitive calcium channel inhibitor GsMTx4 (10 μM) or BT-3033 and GsMTx4 together, which were all added during plating. Graphs show mean % ± SD from 3 independent experiments. *P = 0.05, **P < 0.01, **** P < 0.001, RM-one-way ANOVA with Holm-Šídák’s multiple comparisons test comparing the Sca-1^high^ populations. (H) RT-qPCR analysis of CRC organoids embedded in Matrigel or Collagen I, with addition of DMSO, the Piezo-activator Yoda1, YAP1/TEAD inhibitor K-975 or Yoda1 together with K-975. Graph shows mean expression ± SD from at least 3 independent experiments. ***p < 0.005, ****p < 0.001; RM-one-way ANOVA with Holm-Šídák’s multiple comparisons test performed comparing the ΔΔCT values per gene. (I) Quantification of cells with low, mid or high Sca-1 expression by flow cytometry of cells derived from CRC organoids embedded in Matrigel (M) or Collagen I (C) for 1 day, either in presence of DMSO, the Piezo activator Yoda1 (5 μM), or Yoda1 together with the YAP1/TEAD inhibitor K-975 (10 μM), which were all added during plating. The graph shows mean % ± SD from 3 independent experiments. *P < 0.05, **** P < 0.001, RM-one-way ANOVA with Holm-Šídák’s multiple comparisons test comparing the Sca-1^high^ populations. (J) Representative images of CRC organoids embedded in Matrigel for 1 day in the presence of DMSO or Yoda1 (5 μM) that were added during plating, and immunostained for Sca-1. All scale bar represent 50 μm.

## Discussion

This work uncovers how mechanical signals originating from tumor-ECM interactions control cell fate transitions at the metastatic onset of CRC. We propose that the dissemination of CRC cells from the primary tumor is coupled to their reversion into a fetal-like state, mediated by integrins and mechanosensitive calcium channels that respond to pulling of disseminating cells on the collagen I-rich interstitial matrix.

The fetal-like state is maintained as CRC cells enter the circulation and enables the seeding of metastases, thereby explaining recent findings that identified Lgr5^neg^ cells as the predominant metastasis-initiating cell population of CRC (Fumagalli et al., 2020; Heinz et al., 2022). Whereas fetal-like cells can drive the initial phase of metastasis formation, the eventual re-emergence of Lgr5^pos^ CSCs is essential for the overt outgrowth into macro-metastasis and colonization of the host organ (Cheung et al., 2020; Heinz et al., 2022). This implies that fetal-like cells and Lgr5^pos^ CSCs both possess stem-cell potential, but function in different phases of metastasis formation. Which distinct characteristics of CSCs and fetal-like cells underly their differential roles, and what drives the reappearance of Lgr5^pos^ CSCs after the initial outgrowth of a secondary tumor, warrants further investigations.

While reversion of intestinal cells into a fetal-like state was first linked to colitis-associated regeneration (Yui et al., 2017), from this and recent other studies it emerges that cellular reprogramming into this state contributes to intestinal tumorigenesis. Acquisition of fetal intestinal identity has recently been linked to tumor initiation in both an APC^Min/+^ and right-sided CRC mouse model, downstream of biochemical signals originating from intestinal fibroblasts and microbial-driven inflammation, respectively (Roulis et al., 2020; Leach et al., 2021). The fetal-like state has further been associated with resistance of CRC cells to inhibition of Wnt signaling and to chemotherapy (Han et al., 2020; Solé et al., 2022; Vasquez et al., 2022). Our findings advance on these previous studies by showing that CRC reversion into a fetal-like state is intrinsically linked to the metastatic cascade, being induced by interactions of disseminating cells with the Collagen I-rich matrix and promoting their metastatic potential. Furthermore, our organoid model demonstrates that activation of mechanosensitive calcium channels associated with pulling on the collagen network suffices for the fetal-like reprogramming of CRC cells. Future studies may reveal whether additional factors previously implicated in this reprogramming (e.g. TGFβ (Han et al., 2020) and inflammation (Leach et al., 2021)) contribute to fate transitions during the metastatic cascade. Conversely, mechanical signals transduced by integrins and mechanosensitive calcium channels could potentially fulfill a broader role in controlling cellular plasticity during other steps of CRC tumorigenesis.

Although our findings associate disseminating and metastatic CRC cells with a fetal-like program, recent reports have identified alternative gene expression programs marking CRC cells with high metastatic potential (Ganesh et al., 2020; Sacchetti et al., 2021; Cañellas-Socias et al., 2022). In part, these programs are also specific to Lgr5^neg^ cells, including the cell population marked by EMP1 expression that is responsible for metastatic relapse following removal of the primary tumor (Cañellas-Socias et al., 2022). Other markers define metastatic CRC cell populations irrespective of Lgr5 status, including expression of L1CAM that was shown to be induced by loss of neighbor cell contact during tumor cell dissemination (Ganesh et al., 2020). The lack of universal molecular characteristics across these identified cell populations supports the idea that the metastatic-initiating potential can be acquired by different molecular traits that introduce stem-like properties (Massagué and Ganesh, 2021). Mutational and tumor-environmental factors may determine which program is utilized. For instance, stromal-rich CMS4 subtype tumors could rely mostly on fetal-like reprogramming (Vasquez et al., 2022).

Integrin receptors are well-recognized to transmit mechanical information to YAP1 signaling (Dupont, 2016). Our data support this role of integrins in the YAP1-dependent reprogramming of CRC cells by matrix-associated pulling forces, but reveal that integrins act in concert with mechanosensitive calcium channels in this process. Our data implicate the mechanosensitive channel Piezo1 in this, as its chemical activation is sufficient to induce the fetal-like state in CRC organoids, yet other mechanosensitive channels expressed in CRCs (including channels of the Trp family) may partake in this. Piezo and Trp channels were recently shown to be activated at sites associated with high pulling forces on the ECM in 2D-cell culture models (Ellefsen et al., 2019; Yao et al., 2022; Kerstein et al., 2013; Wei et al., 2009) and physically interact with integrin-based focal adhesions (Chen et al., 2018; Yao et al., 2022; McHugh et al., 2010). As CRC cells protrude and exert tensile forces on the Collagen I-rich matrix, the local build-up of tension could thus promote opening of mechanosensitive channels, which is supported by our observation of elevated pulses of calcium at sites of aligned collagen fibers. The mechanism linking calcium channel activation to YAP1-driven transcription remains to be identified, and both Hippo-dependent and independent mechanisms have been proposed to link mechanosensitive calcium influx to YAP1 (Pathak et al., 2014; Sharma et al., 2019; Kim et al., 2022; Duchemin et al., 2019; Zhou et al., 2020; Liu et al., 2021). Similarly, the interplay between integrins and mechanosensitive calcium channels in CRCs remains to be resolved, and moreover, these mechanosensors are described to reciprocally affect each other (Swaminathan and Gloerich, 2021; De Felice and Alaimo, 2020). As such, local influx of calcium can increase myosin contractility and impact focal adhesions dynamics (Pardo-Pastor et al., 2018; Chen et al., 2018; Yao et al., 2022; Yu and Liao, 2021), thereby providing a potential feedback mechanism by which calcium channel opening by local pulling forces could reinforce their own stimulation.

Altogether, our findings shed light on the regulation of cell fate transitions underlying CRC metastasis and provide novel insights into the role of mechanosensitive calcium channels in tumor malignancy. Future studies may delineate whether other pro-metastatic processes influenced by calcium influx contribute to the mechanical regulation of CRC progression (Monteith et al., 2017), and how integrins and mechanosensitive channels interact with other force-sensitive mechanisms that may impact intestinal tumors (Brás et al., 2022; Broders-Bondon et al., 2018).

## Methods

### Antibodies and reagents

The following commercial antibodies were used at the indicated concentrations for immunofluorescence (IF), immunohistochemistry (IHC) and flow-cytometry (FC): Pacific Blue anti-mouse Ly-6A/E (Biolegend, Cat#108120, 1:50 FC), Sca-1/Ly6A/E (E13 161-7, Abcam, Cat#ab51317, 1:200 IF), YAP1 (63.7, Santa Cruz, Cat#sc-101199, 1:200 IF), Annexin A1 (Cell Signaling, CS32934, 1:400 IHC), ANXA1 (Sigma, HPA011271, 1:100 IF of 3D grown organoids, 1:500 IF of liver tissue), Epcam (Cell Signaling, CS32934, 1:200 IHC), GFP (Abcam, ab6673, 1:1000 IF liver slices), RFP (Rockland, 600-401-379, 1:300 IF liver tissue) and Phospho-FAK (Tyr397) (Invitrogen, Cat#44-624G, 1:100 IF). The following reagents were used at the indicated concentrations: 100 μg/mL doxycycline (Bio-Connect), 10 μM K-975 (MedChemExpress), 5 μM Yoda 1 (Tocris), 10 μM Y27632 (Gentaur), 10 μM ML-7 (Tocris), 5 μM BTT 3033 (Tocris) and 10 μM GsMTx4 (My BioSource).

### Plasmids

For forced expression of constitutively active YAP1 (YAP1-5SA), Myc-YAP1-5SA from the pQCXIH-Myc-YAP1-5SA plasmid (Addgene #33093) was recloned in a pDONR221 backbone and the Myc-tag was replaced with a Flag-tag (Epoch Life Science). The sequence 5’-CAATGCGGAATATCAATCCCAGCACAGCAAATTCTCCAAAATGTCAGG-3’ was added between base pairs 982 and 983 of the YAP1 gene, similar to the previously described YAP1 constructs (Panciera et al., 2016). The Flag-YAP1-5SA construct was inserted into a pInducer20 backbone with blasticidine resistance cassette to generate pInducer20-FLAG-YAP1-S5A-Ubc-rtTA-IRES-Blast x pLV. The pLV-K-GECO1-IRES-Puro plasmid was generated by In-Fusion cloning of the insert of the original pcDNA-K-GECO1 vector (Addgene #105865; Shen *et al*., BMC Biol 2018) into a lentiviral pLV vector.

### Organoid cultures

Colorectal cancer organoids isolated from tumorigenic colons of Villin^Cre-ERT2^; APC^fl/fl^; KRAS^LSL-G12D/+^; TP53^KO/KO^; R26R-Confetti; Lgr5^DTR-eGFP^ transgenic mice (Fumagalli et al., 2020) were cultured in matrigel (Corning) drops in culture medium containing advanced DMEM/F12 advanced medium (Life Technologies), 10 mM HEPES (Life Technologies), 1x Glutamax (Life Technologies), 100 Units/mL Penicilin-Streptomycin (Westburg), 10% Noggin-conditioned medium, 2% B27 (Thermo Fisher), 1.25 mM N-Acetyl-L-Cysteine (Sigma) and 100 μg/mL Primocin (Bio-Connect), and passaged every 3 or 4 days. In experiments with GsMTx4 treatment, N-Acetyl-L-Cysteine was omitted from the culture medium. Patient-derived P16T colorectal cancer organoids (Van de Wetering et al., 2015) were cultured in Basement Membrane Extract (BME, R&D systems) drops in culture medium containing advanced DMEM/F12 medium containing 10 mM HEPES (Life Technologies), 1x Glutamax (Life Technologies), 100 Units/mL Penicilin-Streptomycin (Westburg), 1x B27 (Fisher Scientific), 1.25 mM N-acetyl-cysteine (Sigma), 10 mM Nicotinamide (Sigma), 3 μM SB202190 (Gentaur), 500 nM A83 (Bio-Techne) and 50 ng/ml stock EGF (Peptrotech) and passaged every 7 days. Organoids lacking Confetti-Tdimer2 (RFP) expression were generated by CRISPR/Cas9-based gene editing. The pSpCas9(BB)-2A-Puro (PX459) V2.0 plasmid (Addgene #62988) containing a tdimer2-specific gRNA (guide: 5’-CGGCCACGAGTTCGAGATCGAGG-3’) was transfected in the organoids by electroporation. Knockout cells were selected with flow cytometry for absence of tDimer2 expression and grown as a polyclonal line. Organoids expressing the K-GECO1 calcium sensor were generated by lentiviral transduction of organoids lacking Tdimer2 expression with the pLV-K-GECO1-IRES-Puro plasmid. To generate lentiviral particles, HEK293T cells were transfected with the lentiviral plasmids together with 3^rd^ generation packaging vectors. Lentiviral particles were concentrated using the LentiX Concentrator (Clontech). Organoids were trypsinized into small cell clusters and incubated with the concentrated virus particles supplemented with 4 μg/mL Polybrene (Sigma Aldrich), for 60 minutes while centrifuging at 600 rpm at RT followed by 4 hr incubation at 37 °C. The organoids were plated in Matrigel and selection for transduced organoids was performed 3 days after infection by supplementing the organoid culture media with 20 ug/mL blasticidin (Cat# ant-bl-05, Invitrogen).

### 3D matrigel and collagen I assays

Intact organoids were isolated from matrigel cultures using 1 mg/mL Dispase II (Cat# 17105041, Life Technologies) or by mechanical dissociation of the matrigel drops followed by 3 washes with DMEM/F12 media. Organoids were either embedded in Matrigel or in type I Collagen lattices consisting of non-pepsinized rat-tail Collagen at a final concentration of 1 mg/ml. The collagen mixture was prepared in buffer containing 10X phosphate buffer saline (PBS) (Gibco), NaOH and dH2O according to the manufacturer’s instructions and the collagen I gel was pre-polymerized on ice for 2-3 hours, after which organoids were embedded in the collagen I gel followed by incubation at room temperature for 5 minutes. The collagen I- and Matrigel gels were plated as 45 μL drops on a pre-warmed 24-well culture plate (Costar). The plate was incubated at 37 °C until polymerization of the gels was achieved, after which the culture medium was added. Perturbation studies were performed with corresponding DMSO or MQ controls. All analyses were performed ∼24h following plating of the organoids.

### Single cell RNA sequencing

Organoids cultured in Matrigel or Collagen I for 1 day were isolated from the gels using 20 mg/mL collagenase (Sigma) in PBS for 3 minutes at 37 °C, washed in PBS and dissociated into single cells by TrypLE treatment (Fisher Scientific) at 37 °C. Cells were washed in PBS containing 1% FCS and 1 mM EDTA, subsequently filtered through a Filcon 50 uM stainer (BD BioSciences), and plated as single cells in 384-well plates by FACS (FACS Aria III, BD Biosciences). Single-cell mRNA-sequencing was performed by Single Cell Discoveries (Utrecht, the Netherlands) according to the Sort-seq protocol (Muraro et al., 2016). Libraries were sequenced on an Illumina NextSeq500 at paired-end 60-and 26-bp read length and 150.000 reads per cell. Reads were mapped to the mm10 RefSeq genome. Analysis was performed using the Seurat R-package (v 4.3.0)(Stuart et al., 2019). Cells with at least 10.000 total transcripts were included in the analysis. The first 16 principle components were used for unsupervised hierarchical clustering (default settings at resolution 0.5). Differential gene expression was performed using a Wilcoxon Rank Sum test (FindAllMarkers with min.pct = 0.1). Genes were ranked by p-value and the top 105 genes per group were displayed (minimal p-value adjusted = 3.16e-13). Scores for gene signature expression were calculated using the AddModuleScore function.

Of previously published single-cell RNA sequencing data (Fumagalli et al., 2020) cells with at least 1000 total transcripts were included in the analysi.

### GSEA analysis

Gene set enrichment analysis (GSEA) was performed using the Broad Institute GSEA tool (software.broadinstitute.org/gsea/index.jsp, v 4.2.3) with default settings. HALLMARK genesets were derived from the Molecular Signatures Database (Liberzon et al., 2015). Further comparisons were made with gene sets for intestinal cell types (Haber et al., 2017), a YAP1 overexpression gene signature (Gregorieff et al., 2015), a fetal spheroid signature (Mustata et al., 2013) and a DSS-induced intestinal colitis signature (Yui et al., 2017). Results with a FDR q value less than 0.25 and p-value less than 0.05 are reported.

### RT-qPCR

Matrigel-and collagen I plated organoids were incubated with 20 mg/mL collagenase (Sigma) in PBS for 3 minutes at 37 °C, washed once with cold PBS, and subsequently lysed in RLT buffer (Qiagen). Total RNA was isolated using the RNeasy kit (Qiagen) with DNase treatment (Qiagen), and cDNA was synthesized using the iScript cDNA Synthesis Kit (Bio-Rad). RT-qPCR was performed with FastStart SYBR Green Master mix (Roche) with optimized primer pairs. The fold changes were determined using the ΔCt method with normalization to the pooled expression of housekeeping genes (*Pbgd, Tuba, Hnrnpa* and *Ppia*). The following primers were used:

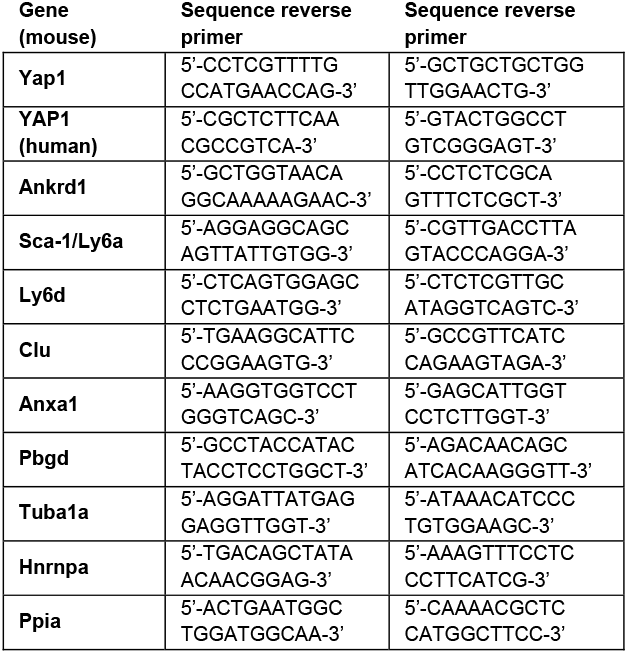

### Immunofluorescence of 3D-cultured organoids

Organoids in Matrigel and Collagen I were fixed with 4% paraformaldehyde for 10 minutes at room temperature, washed 3 times with PBS and blocked in PBS containing 10% Normal Goat Serum (NGS) and 0.3% Triton X-100. Samples were incubated with the primary antibodies (diluted in 1% BSA, 0.3% Triton-X100 in PBS) overnight at 4 °C while shaking, after which the samples were washed 4 times with PBS for 10 minutes. The samples were incubated with Alexa-conjugated secondary antibodies (1:500; Life Technologies) and DAPI (5 μg/mL, Sigma) overnight at 4 °C while shaking and subsequently washed 4 times with PBS for 10 minutes. The samples were imaged using a SP8 confocal microscope (Leica Microsystems, x water objective, N.A. = 1.1) or LSM880 confocal microscope (Zeiss, 40x water objective, N.A. = 1.1). Confocal reflection microscopy was used to visualize the Collagen I fibres. The samples were illuminated with a 488 nm Argon laser and the reflected light was detected in the 480-500 nm range. For analysis of the fraction of Lgr5^pos^ cells or cells with nuclear YAP1 per organoid, cells with positive Lgr5-eGFP signal or nuclear YAP1 intensity above cytosolic YAP1 intensity were counted in a single plane in the middle of each organoid and divided over the total number of cells in that plane.

### Live-cell microscopy

Brightfield imaging and real-time analysis of Lgr5-eGFP expression of organoids 1 day after plating in Matrigel or Collagen I gels were performed on a Nikon Spinning Disc confocal microscope using a 40X water objective (NA = 1.15) in a temperature-and CO_2_-controlled incubator, using NIS-Elements software with a 15 minute time interval. To monitor intracellular calcium levels, organoids expressing the K-GECO1 sensor embedded in Matrigel or Collagen I in 96-well imaging plates (Cellstar) were imaged on the same microscope with 5 seconds interval. For analyses of calcium peaks, we generated single cell traces (15 minutes long) of intensity of the K-GECO1 sensor using ImageJ software and identified peaks using the BAR plugin “Find Peaks” (Ferreira et al., 2015). To determine peak amplitude, we divided the maximum intensity of each peak (Fmax) by the baseline intensity of the sensor (10% lowest values per cell, F0). Peak frequency was determined by calculating the number of peaks with Fmax/F0 ≥ 1.4.

### 2D culture of organoids

Polyacrylamyde gels were fabricated on glass-bottom dishes (World Orecision Instruments). First, the glass was treated with silane (3-(Trimethoxysilyl) propyl methacrylate) in PBS (1:3) for 15 minutes. After 3 washes with water, the glass was treated with 0.5% glutaraldehyde in PBS for 30 minutes. After thorough washing and drying, a 20 μl drop of gel mix was polymerized between the glass and a 18 mm coverslip. Depending on the stiffness, the gel mix was made as follows:

| Stiffness | PBS | Acryl-<br>amide<br>40% | Bis-<br>acryl-<br>amide<br>2% | APS | TEMED |
| --- | --- | --- | --- | --- | --- |
| 0,5 kPa | 439,75 | 50 | 7,5 | 2,5 | 0,25 |
| 5 kPa | 392,95 | 93,3 | 11 | 2,5 | 0,25 |
| 30 kPa | 309,75 | 150 | 37,5 | 2,5 | 0,25 |

After 1 hour of polymerization, gels were incubated in PBS and the glass coverslip was removed with a scalpel. Gels were then activated with 75ul of Sulpho-SANPAH 2ug/mL incubated under 365 UV light for 7.5 minutes. Gels were washed twice with 10mM HEPES for 3 minutes under agitation, and once with PBS, before incubating with coating solution overnight at 4°C. To compare Matrigel and Collagen I, the coating was 10% Matrigel (corning) or 10 μg/ml of rat tail collagen I (Corning) in PBS. To compare different stiffnesses, 1 μg/mL of collagen was used.

The day after coating, gels were washed once with PBS and let dry for 5 minutes before organoid seeding. Organoids were dissagregated into single cells in two steps, first mechanically with a 18G needle, then by incubating with TrypLE for 5 minutes. 10.000 cells were seeded on each gel. Cells were fixed after 24 hours of culture using 4% PFA for 15 minutes and mounted in Aqua-Poly/Mount. Imaging was performed with a Spinning disk microscope (CSU-W1 Yokogawa) using a 40x objective (Water Immersion, NA 1.15).

### Flow cytometry

Organoids cultured in Matrigel or Collagen I for 1 day were isolated from the gels using collagenase (20 mg/mL) for 3 minutes at 37 °C, washed once with cold PBS, and dissociated into single cells by TrypLE treatment (Life Technologies) at 37 °C. Following washing with PBS containing 1 mM EDTA and 1% FCS, cells were fixed with 4% PFA in PBS and incubated with primary antibody (in 1% BSA in PBS) for 20 minutes at room temperature. Samples were washed with PBS and flow cytometry was performed using the BD FACS Celesta. For analysis, individual experiments were normalized by scaling the Lgr5 expression (excluding the 2.5% lowest and highest expressors) to a 0-100 range. Thresholds for binning into low/high expression were <10 / >30 and <10 / >40 for Lgr5 and Sca-1, respectively (see Figs S1c and s2d for further clarification of these thresholds).

### *In vitro* outgrowth assay

Organoids cultured in Matrigel or Collagen I for 1 day were isolated from the gels using collagenase (20 mg/mL) for 3 minutes at at 37 °C, washed once with cold PBS, and dissociated into single cells by TrypLE treatment (Life Technologies) at 37 °C. Following washing with PBS containing 1 mM EDTA and 1% FCS, cells were filtered through a Filcon 50 μM strainer (BD BioSciences) and incubated with Pacific Blue anti-mouse Ly-6A/E antibody (Biolegend). Cells were washed with PBS containing 1 mM EDTA and 1% FCS, resuspended in culture medium supplemented with Y27632 (10 μM, Gentaur) and DRAQ7 (1.5 μM, Cell Signaling), sorted based on Lgr5 and Sca-1 expression (and absence of DRAQ7) using the FACS Aria III (BD Biosciences). Sorted cells were embedded as single cells in Collagen I gels at a density of 200 cells/μl in 96-well imaging plates (Cellstar) and gels were supplemented with organoid culture medium supplemented with DRAQ7 (3 μM). Cells were imaged on a Cell Observer microscope (Zeiss) using a 20x dry objective (N.A. = 0.75) in a temperature and CO_2_-controlled incubator with a 4 hour time interval. Cells/organoids were identified by the expression of Confetti-tDimer2 (RFP), with dead cells being identified by staining for DRAQ7 (Cell Signaling).

### Mesenteric vein injection

All mouse experiments were approved by animal experimentation committee at the Netherlands Cancer Institute (NKI-AVL) and performed according to institutional and national guidelines. Mice were housed in temperature-controlled facilities (20–24 °C) at 40–70% humidity with a 12 h dark/light cycle and *ad libitum* access to food and water. Six week old male NOD-SCID mice were purchased from Charles River Laboratories and were randomly assigned to experimental groups. 50.000 FACS-sorted cells were resuspended in 100 μl of sterile PBS and injected in the mesenteric vein of acceptor mice as previously described (Fumagalli et al., 2017; Van Der Bij et al., 2010). Mice were sedated with isoflurane inhalation anesthesia (∼2% isoflurane/ O2 mixture). Before and post-surgery, the mice were treated with a sub-cutaneous dose of buprenorphine (Buprecare, Multidosis-Astfarma, 3mg per mouse). Animals were sacrificed 30 days post-injection. Livers and lungs were fixed periodate-lysine-4% paraformaldehyde (PLP) buffer (McLean and Nakane, 1974) overnight at 4 °C, incubated in 30% sucrose overnight at 4°C, embedded in Tissue-Tek (Sakura, cat. no. 4583) and stored at -80°C. Whole livers were sectioned in 30 μm thick slices and liver slices were imaged on a Cell Observer microscope (Zeiss) using a 5x dry objective (N.A. = 0.16). The number of metastases were quantified using ImageJ-and Zen blue Software (Zeiss). Quantification was performed in a blind manner. For immunofluorescence imaging of the liver tissue, slices were rehydrated in PBS and blocked in PBS containing 5% NGS and 0.5% Triton X-100. Slices were incubated with primary antibodies (diluted in 5% NGS and 0.1% Triton X-100) overnight at 4 °C, after which the slices were washed 3 times with 0.1% Triton X-100 in PBS and 1 time with PBS. The slices were incubated with Alexa-conjugated secondary antibdoies (1:500; Life Technologies) and DAPI (5 μg/mL, Sigma) for 1 hour at RT and subsequently washed 3 times with 0.1% Triton X-100 in PBS and 1 time with PBS. Sections were embedded in Vectashield Plus Antivade mounting medium (Vectorlabs) and imaged using the LSM880 confocal microscope (Zeiss, 40x water objective, N.A. = 1.1).

### Immunohistochemical analysis of patient tissue

Formalin fixed paraffin embedded (FFPE) patient samples were obtained from a prospective HIPEC database, containing tumor samples of patients with peritoneal metastasized colorectal cancer. Sections were deparaffinized and incubated with 1.5% hydrogen peroxide solution for 20 minutes. Next, slides were boiled in a citrate antigen retrieval buffer (pH 6). Sections were incubated overnight at 4 °C. Bound antibodies were detected by BrightVision poly-horseradish peroxidase conjugated goat anti-rabbit secondary antibo y (Immu ologic, VWRKDPVR110HRP). The sections were then developed with diaminobenzidine (DAB) at RT for 10 minutes and counterstained using hematoxylin.

### Quantification and statistical analysis

Statistical analysis was performed using Graphpad prism 9. Gaussian distribution of the data was tested by a Shapiro-Wilk test to next apply parametric or non-parametric statistics. Details of the performed statistics, n values and p values are described in the figure legends.

## Acknowledgements

We thank Willem-Jan Pannekoek (UMC Utrecht) for critical reading of the manuscript, Ingrid Jordens and Susan Zwakenberg (UMC Utrecht) for assisting with the FACS sort experiments, Arianna Fumagalli and Lennart Kester (Netherlands Cancer Institute) for providing reagents and datasets, Single Cell Discoveries (Utrecht, The Netherlands) for performing single cell RNA sequencing, Koen Oost, Maria Heinz, Johan de Rooij (UMC Utrecht) and members of our laboratory for helpful discussions throughout the project and Joep Sprangers and Susanna Plugge (UMC Utrecht) for help with analysis of the scRNAseq data. This work was supported by the Dutch Cancer Foundation (KWF-12345) and the Netherlands Organization for Scientific Research (NWO; 016.Vidi.189.166, NWO gravitational program CancerGenomiCs.nl 024.001.028, and the Science-XL research program The Active Matter Physics of Collective Metastasis 2019.022).

## Supplementary figures

**Figure S1.**
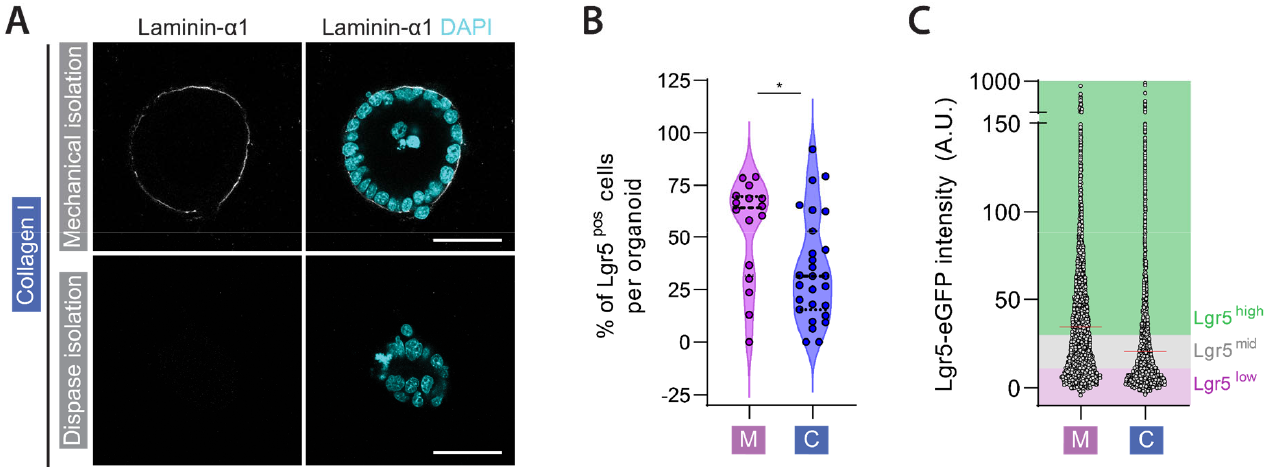
Characterization of CRC organoids in Matrigel and Collagen I. (A) Representative images of CRC organoids embedded in Collagen I following isolation with dispase treatment compared to mechanical isolation, and immunostained for Laminin-α1 (a main component of Matrigel and basement membrane) after 1 day. Scale bar represents 50 μm. (B) Quantification of the percentage of Lgr5^pos^ cells per organoid, in organoids embedded in Matrigel or Collagen I for 1 day. Every data point represent a single organoid, data were pooled from 2 independent experiments with at least 8 organoids per condition in each experiment. Black bars represent mean ± quartiles. *P < 0.05, Mann-Whitney test. (C) Analysis of Lgr5-eGFP expression by flow cytometry of cells from CRC organoids embedded in Matrigel (M) or Collagen I (C) for 1 day, showing the thresholds of Lgr5 expression used to bin cells into Lgr5^low^, Lgr5^mid^ and Lgr5^high^ populations. Every data point represents individual cell, data were pooled from 3 independent experiments.

**Figure S2.**
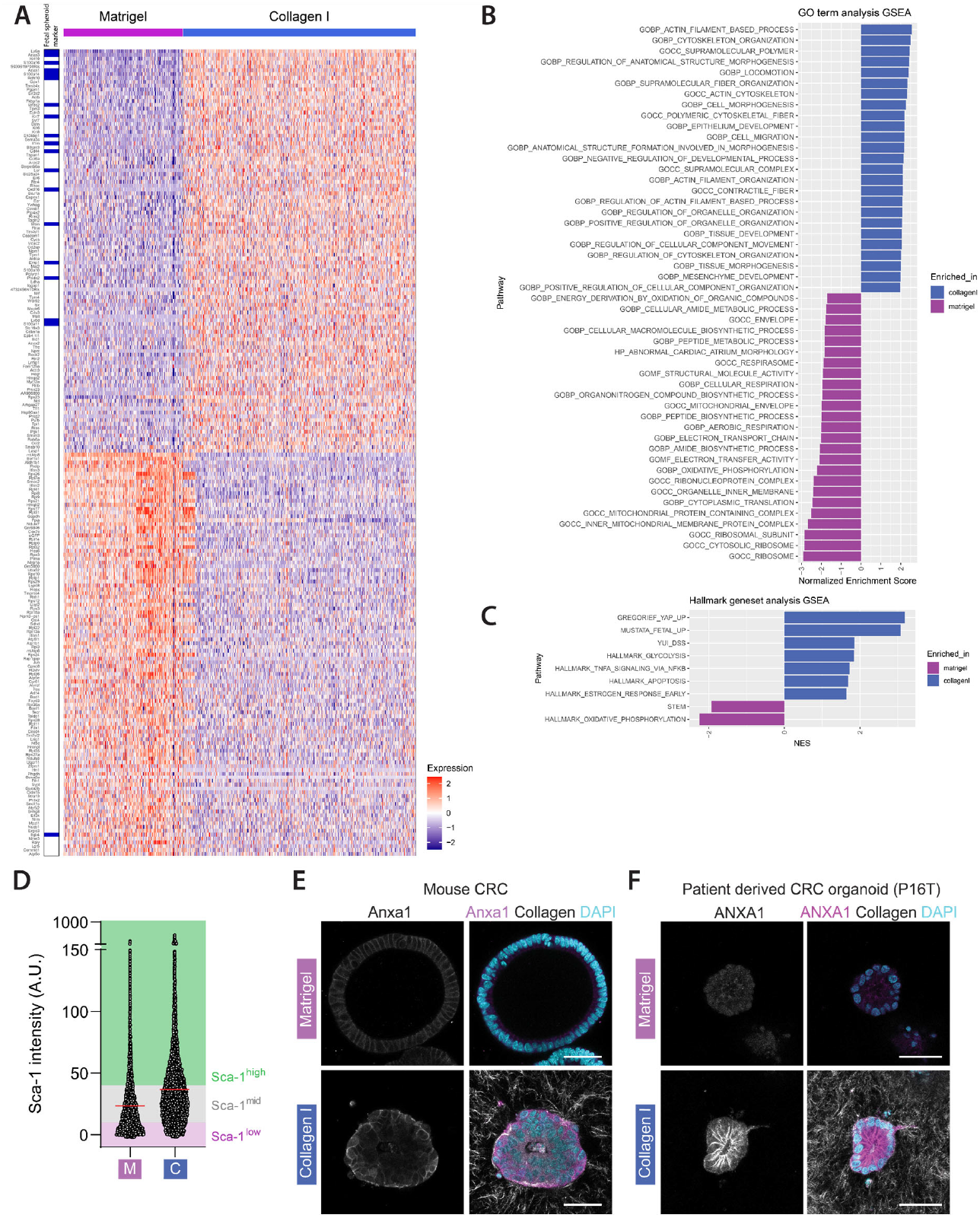
Expression analyses of CRC organoids in Matrigel and Collagen I. (A) Heatmap showing scaled expression values for top 105 markers for each group (Matrigel or Collagen I). Genes highlighted in blue are part of the fetal spheroid gene signature (Mustata et al., 2013). (B) GO-term analysis comparing expression profiles of cells originating from Matrigel or Collagen I embedded organoids, showing top 25 associated GO-terms for each condition. (C) GSEA analysis comparing expression profiles of cells originating from Matrigel or Collagen I embedded organoids. Comparisons were made to HALLMARK gene sets, genesets for intestinal cell types (Haber et al., 2017), YAP1 overexpression gene signature (Gregorieff et al., 2015), fetal-spheroid gene signature (Mustata et al., 2013), and DSS-induced intestinal colitis signature (Yui et al., 2017). (D) Analysis of Sca-1 expression by flow cytometry of cells from CRC organoids embedded in Matrigel (M) or Collagen I (C) for 1 day, showing the thresholds of Sca-1 expression used to bin cells into Sca-1^low^, Sca-1^mid^ and Sca-1^high^ populations. Every data point represents individual cell, data were pooled from 3 independent experiments. (E, F) Representative images of mouse CRC organoids (E) and human patient-derived CRC organoids (P16T, (van de Wetering et al., 2015)) (F) embedded in Matrigel or Collagen I for 1 day and immunostained for AnnexinA1 (ANXA1). Collagen I fibers are visualized by reflection microscopy. Scale bars represent 50 μm.

**Figure S3.**
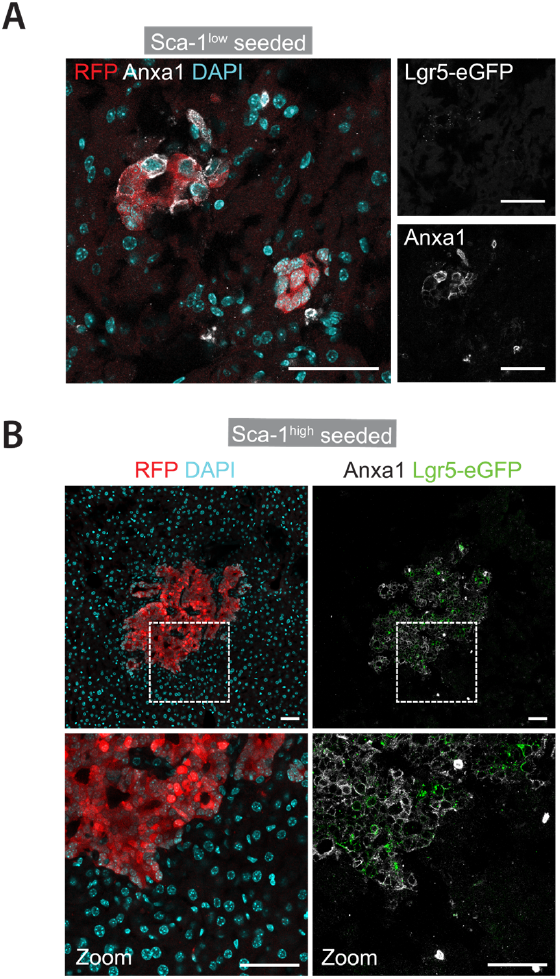
Re-expression of Sca-1 expression in Sca-1^neg^ seeded metastasis. (A) Representative image of immunostaining for AnnexinA1 (ANXA1) of a liver slice showing ANXA1^pos^ metastatic foci originating from Sca-1^low^ injected single cells. (B) Representative image of immunostaining for AnnexinA1 (ANXA1) and Lgr5-eGFP expression marking cancer stem cells in a liver slice showing Lgr5-eGFP^pos^ metastatic foci originating from Sca-1^high^ injected single cells. All scalebars represent 50 μm

**Figure S4.**
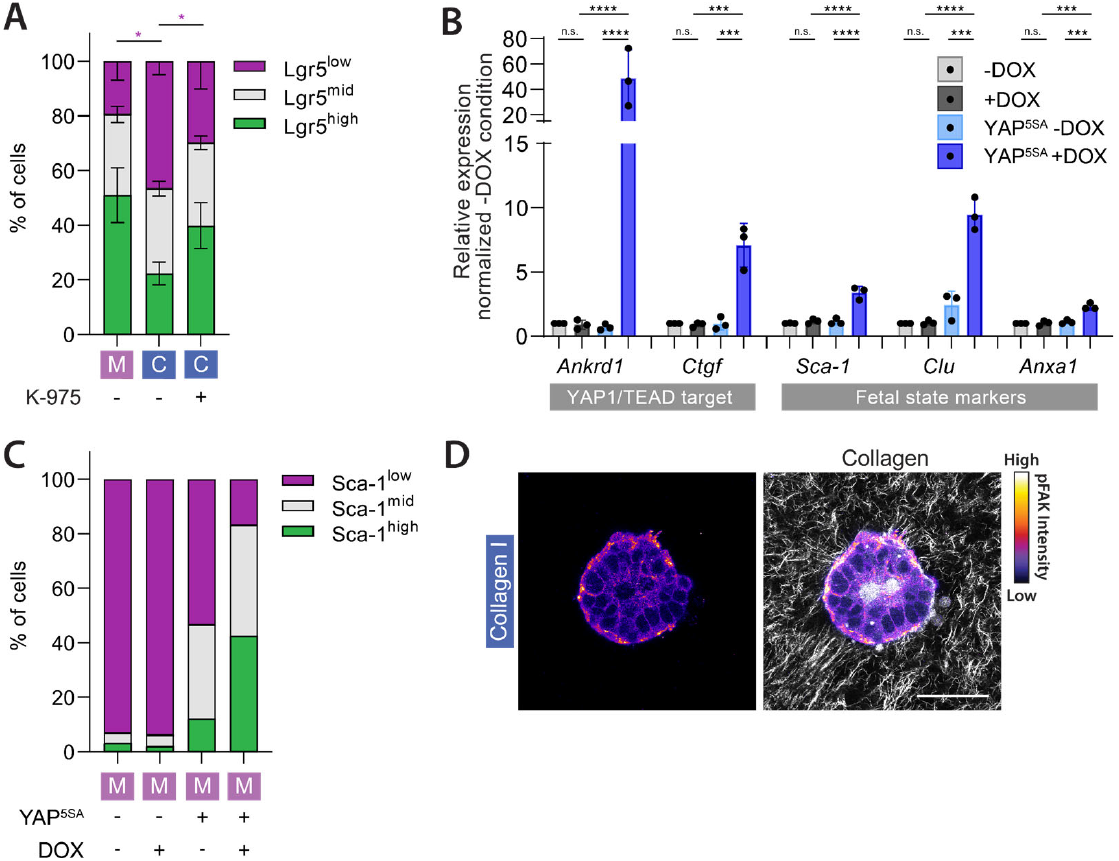
YAP1-dependent reprogramming of CRC cells into fetal-like state. (A) Quantification of cells with low, mid or high Lgr5 expression by flow cytometry of cells derived from CRC organoids embedded in Matrigel (M) or Collagen I (C) for 1 day, with addition of DMSO or the YAP1/TEAD inhibitor K-975 (10 μM) that were added during plating. Graph shows mean % ± SD from 3 independent experiments. *P < 0.05, RM-one-way ANOVA with Holm-Šídák’s multiple comparisons test comparing the Lgr5^low^ populations. (B) RT-qPCR analysis of CRC organoids embedded in Matrigel or Collagen I, with or without doxycycline-inducible YAP1^5SA^ expression for 1 day. Graph shows mean expression ± SD from 3 independent experiments. ***p < 0.005, ****p < 0.001; RM-one-way ANOVA with Holm-Šídák’s multiple comparisons test performed comparing the ΔΔCT values per gene. (C) Quantification of cells with low, mid or high Sca-1 expression by flow cytometry of cells derived from CRC organoids embedded in Matrigel (M) or Collagen I (C), with or without doxycycline-inducible YAP1^5SA^ expression for 1 day. (D) Representative images of CRC organoids embedded in Collagen I and immunostained for FAK pY397 (from Fig. 4f) with Collagen I fibers visualized with reflection microscopy. Scale bar represents 50 μm.

## Supplementary movies

**Movie 1**

Time lapse brightfield imaging of a CRC organoid embedded in Matrigel. Scalebar represents 25 μm. Time indicated in hrs:min.

**Movie 2**

Time lapse brightfield imaging of a CRC organoid embedded in Collagen I. Scalebar represents 25 μm. Time indicated in hrs:min.

**Movie 3**

Time lapse confocal imaging of Matrigel-embedded CRC organoids expressing the K-GECO-1 calcium sensor (Fire). Scalebar represents 25 μm. Time indicated in hrs:min:sec.

**Movie 4**

Time lapse confocal imaging of Collagen I-embedded CRC organoids expressing the K-GECO-1 calcium sensor (Fire). Scalebar represents 25 μm. Time indicated in hrs:min:sec.

